# Long-term phenotypic effects following vitrified-thawed embryo transfer in a rabbit model

**DOI:** 10.1101/410514

**Authors:** Ximo Garcia-Dominguez, David. S. Peñaranda, Guillem Estruch, José Blanca, Victor García-Carpintero, Joaquín Cañizares, Francisco Marco-Jiménez, José Salvador Vicente

## Abstract

Since the first human was conceived through in vitro fertilisation in 1978, over 6.5 million babies have been born by assisted reproductive technologies (ARTs). Although most ART babies and children seem healthy, in recent years several studies have evidenced a potential impact of ARTs on long-term development and health. Herein, we have developed an animal model to determine whether vitrified embryo transfer procedure induces phenotypic changes over the growth performance and in the complementary transcriptomic and proteomic analyses at hepatic level. To this end, 2 populations were developed; vitrified embryos transferred to the surrogate mothers (VT) and naturally conceived animals (NC). After delivery, animals were weighed weekly from 1 to 20 weeks of age. In adulthood, animals were euthanized and organs were harvested and weighed. After that, liver tissue was used to identify changes in the transcriptomic and proteomic profile. At adulthood, VT group showed significant lower body, liver and heart weight. After functional analysis of RNA-Seq data, a subset of 96 differentially expressed transcripts in VT animal were related to alteration in zinc homeostasis, lipid metabolism, and hepatic immune pathways. After proteomic analysis, a subset of 76 differentially expressed proteins also revealed some disturbed metabolic pathways related with the lipid and glycan metabolism, and an impaired oxidative metabolism related to ATP synthesis in the mitochondria. Current findings suggest that progeny derived after transfer of vitrified embryos have long-term consequences on growth rate and vital organs weights in adulthood, correlated with molecular signatures at transcriptomic and proteomic level of hepatic tissue.

## Introduction

Since the first human conceived through *in vitro* fertilisation in 1978, it has been estimated that more than 6.5 million babies born by assisted reproductive technologies (ARTs, [1]). Nevertheless, from the beginning of the application of ARTs, there has been concern about the influence of these technologies on development, and in consequence several epidemiological studies have reported on this issue, associating ARTs with low birth weight, preterm birth, heart disease, hypertension, hyperlipidaemia, insulin resistance and increased risk of type 2 diabetes or adverse neurodevelopmental outcome [1-5]. However, it is difficult to determine in humans whether these effects are really caused by ARTs per se or originate from either genetic abnormalities or risk factors intrinsic to infertile patients [4]. Based on animal models that avoid these confounding factors, analogous effects have been reported, evidencing long-lasting consequences of ARTs such as glucose intolerance, insulin resistance, cardiometabolic disorders, hypertension, behavioural deficits, memory loss, abnormal hepatic and fat metabolomes, placenta dysfunction, body weight and organ weight changes, altered locomotion and shorter lifespan [6-19]. Till now, the vast majority of fertility researchers have been trying to improve the success of ARTs based on apparently healthy babies at home, but only a few are trying to discern whether ARTs leaves a subtle legacy in offspring [20].

In recent years, there has been an increasing trend towards ART cycles that include a cryopreservation procedure that maximises the efficacy of ovarian stimulation cycles in an IVF treatment by allowing storage of the excess embryos and their later use, but also to enable fertility preservation [21, 22]. Hence, cryopreservation of human embryos is currently more important than ever for the cumulative pregnancy rate after IVF [21]. The cryopreservation is a technique in which gametes and embryos are exposed to cryoprotectant solutions and are stored at sub-zero temperatures until needed for use. Although no studies have evidenced genotoxic or teratogenic effects [23], cryopreservation is known to cause extensive damage with marked deleterious effect on embryonic health, compromising its full-term development due to the associated higher rate of pregnancy complications [24, 25]. In addition, offspring present higher risk of high birth weight, high prenatal growth rate (large for gestational age) and hypertensive disorders [26].

Recently, studies carried out under several omics approaches have reported an increased incidence of metabolic activity disorders, cellular stress and changes in developmental potency that varied depending on the in vitro manipulation conditions [27-32]. ART related disorders in mice and rabbit have been evidenced at placental function level by transcriptomic and proteomic studies [17, 33-35]. Furthermore, disturbances in the protein profile of umbilical veins and foetuses derived from ART have been identified, suggesting molecular disorders that can affect later life stages [36, 37]. It has been demonstrated that in vitro manipulation incurs in molecular anomalies beyond parturition, which have been associated with liver weight changes [7], increased fat deposition [16], lower pancreatic weight and impaired functionality [13], endothelial dysfunction, increased stiffness of the vasculature and arterial hypertension [12], heart physiology [38] and functionality [13], behavioural and anxiety level changes [7, 39], muscle physiology [16] and cardiometabolic profile [40]. However, there is a lack of knowledge and understanding around the role of vitrified embryo transfer procedure in adulthood.

To address this issue, we have developed a rabbit model to study the effects of vitrified embryo transfer procedure on body growth and organs weight in adulthood, and hepatic tissue on a global scale in terms of molecular signatures at transcriptomic and proteomic level.

## Materials and Methods

All chemicals, unless otherwise stated, were reagent-grade and purchased from Sigma-Aldrich Química S.A. (Alcobendas, Madrid, Spain). All the experimental procedures used in this study were performed in accordance with Directive 2010/63/EU EEC for animal experiments and reviewed and approved by the Ethical Committee for Experimentation with Animals of the Universitat Politècnica de València, Spain (research code: 2015/VSC/PEA/00061).

### Animals

Healthy animals used as control (naturally conceived animals), embryo donors and surrogate mothers (vitrified and transferred animals) belonged to the synthetic strains selected by Instituto de Ciencia y Tecnología Animal at the Universitat Politècnica de València since the 80s.

### Experimental Design

Figure 1 illustrates the experimental design. Initially, 2 experimental populations were developed: one from vitrified embryos transferred to the surrogate mothers (VT) and other from naturally conceived animals (NC). In both experimental groups, females were artificially inseminated (AI) with semen of unrelated males from the same strain. In the VT group, 3 days after AI, the embryos were recovered, vitrified and then transferred to surrogate mothers by laparoscopy. Meanwhile, NC offspring were generated letting the females give birth after AI. After delivery, male offspring were weighed every week until adulthood. At 56 weeks of age, animals were euthanised and organs were weighed. From the liver, 6 individual biopsies were obtained (3 VT and 3 NC). Over the same samples, complementary RNA sequencing and proteomic studies were performed.

**Fig 1.**
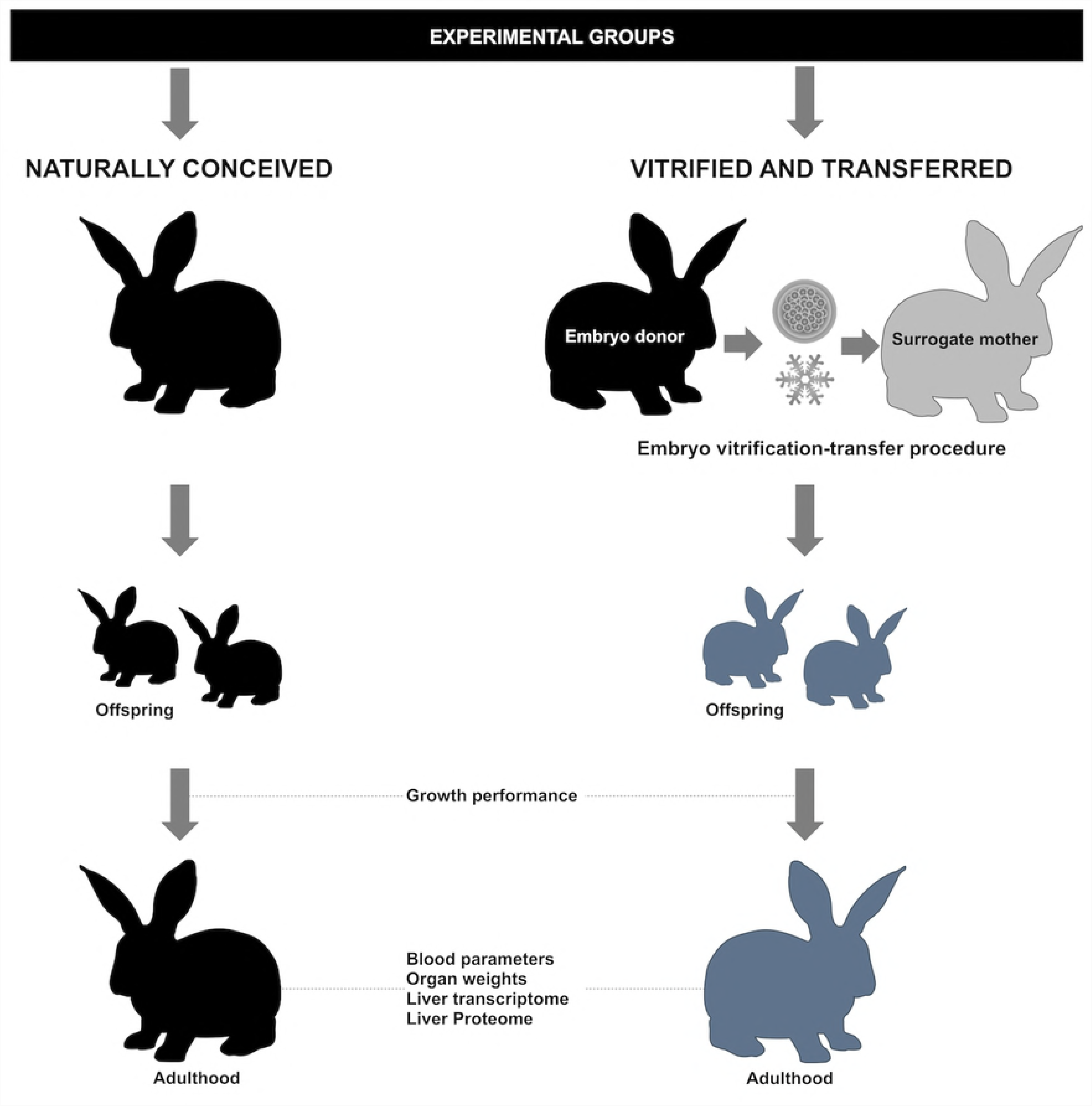
Experimental design. Two experimental groups were developed; (i) animals born from vitrified embryos transferred to the surrogate mothers and (ii) animals born after natural conception. In both groups and after delivery, male offspring were weighed every week until the adulthood (growth performance). At adulthood, haematological and organs weight comparison was performed. In addition, transcriptomic and proteomic comparative analysis of liver tissue was developed.

### In vivo embryo production and collection

A total of 28 donors were AI with semen from mature tested males. When the AI was performed, females were injected with 1 µg of buserelin acetate to induce ovulation. Then, donors were euthanised 72 h post-AI and embryos were recovered. Briefly, the reproductive tract was retrieved and each oviduct and uterine horn (the first one third) was perfused with 5 ml of embryo recovery media, consisting of pre-warmed solution (≈20-25 °C) of Dulbecco’s Phosphate-Buffered Saline (DPBS) solution supplemented with CaCl_2_ (0.132 g/L), 0.2% (w/v) of bovine serum albumin (BSA) and antibiotics (penicillin 100 IU/mL, streptomycin 100 µg/mL and amphotericin B 0.25 µg/mL). After recovery, morphologically normal embryos (correct developmental stage, homogenous blastomeres and intact spherical mucin coat and zona pellucida) were selected and vitrified. A total of 301 embryos were vitrified.

### Vitrification and warming procedure

Embryos were vitrified and warmed using the methodology described by Vicente et al. [41]. Embryos were vitrified in two-step addition procedure at room temperature (≈20-25 °C). In the first step, embryos were placed for 2 min in a solution consisting of 10% (v/v) dimethyl sulphoxide (DMSO) and 10% (v/v) ethylene glycol (EG) in DPBS supplemented with 0.2% of BSA. In the second step, embryos were suspended for 30 s in a solution of 20% DMSO and 20% EG in DPBS supplemented with 0.2% of BSA. Then, embryos suspended in vitrification medium were loaded into 0.125 mL plastic straws (French ministraw, IMV, L’Aigle, France) adding 2 sections of DPBS at either end of each straw, separated by air bubbles. Finally, straws were sealed and plunged directly into liquid nitrogen. Warming was done by horizontally placing the straw 10 cm from liquid nitrogen for 20-30 s and when the crystallisation process began, the straws were immersed in a water bath at 20°C for 10-15 s. The vitrification medium was removed rinsing the embryos into a solution containing DPBS with 0.33 M sucrose for 5 min, followed by one bath in a solution of DPBS for another 5 min.

Only non-damaged embryos (intact mucin coat and pellucid zone) were considered to continue with the transfer. From the 301 vitrified embryos, 287 were recovered successfully and 272 (non-damaged embryos) were catalogued as transferable attending to International Embryo Transfer Society classification.

### Embryo transfer

Embryos were transferred into the oviducts of 26 surrogate mothers by laparoscopy, following the procedure previously described by Besenfelder and Brem, [42]. The mean number of transferred embryos per surrogate mothers was 10.5 (ranged from 6 to 19). Briefly, ovulation was induced in the surrogate mothers with an intramuscular dose of 1 µg Buserelin Acetate 68–72 h before transfer. During laparoscopy, surrogate mothers were anaesthetised by an intramuscular injection of xylazine (5 mg/Kg; Bayer AG) followed by an intravenous injection of ketamine hydrochloride (35 mg/Kg; Imalgene, Merial SA, Lyon, France) 5 min later. During laparoscopy, one dose of morphine hydrochloride (3 mg/Kg; Morfina, B. Braun, Barcelona, Spain) was administered intramuscularly. After surgery, animals were treated with antibiotics (4 mg/Kg of gentamycin each 24 h) and analgesics (0.03 mg/kg of buprenorphine hydrochloride each 12 h and 0.2 mg/kg of meloxicam every 24 h; Alvet Escarti S.L. Guadassuar, Spain) for 3 days.

### Control progeny

The control progeny (NC) group were obtained following the common management of rabbit reproduction without embryo vitrification or embryo transfer procedures. Briefly, contemporaneous control offspring were produced using artificial insemination as a reproduction technique. This procedure was carried out using 0.5 mL of diluted fresh semen from fertile males. Immediately after that, ovulation was induced in inseminated females by an intramuscular injection of 1 µg of buserelin acetate.

### Body growth, organs weight and peripheral blood parameters study

A total of 65 males were weighed weekly from 1 to 20 weeks of age (30 from VT and 35 from NC groups). Then, body growth was estimated by nonlinear regression using the Gompertz curve equation, well suited for rabbits [43]: y= a exp[-b exp(-kt)], where y is the observed body weight of one individual at a specific age (t). The rest of the parameters (a, b and k) of the Gompertz function have a biological interpretation: a can be interpreted as the mature body weight (BW), maintained independently of short-term fluctuations; b is a timescale parameter related to the initial body weight; k is a parameter related to the rate of maturing (growth rate). Furthermore, BW differences between the experimental groups were evaluated weekly. In addition, males were euthanised at week 56 (late adulthood), when the growth plate is closed [44] determining the body weight and vital organs (liver, lungs, heart, kidneys and adrenal glands), spleen and gonads weight.

Prior to euthanasia, individual blood samples were obtained and dispensed into a EDTA-coated tube (Deltalab S.L., Barcelona, Spain). Within 10 minutes of collection, samples were analysed using an automated veterinary haematology analyser MS 4e automated cell counter (MeletSchloesing Laboratories, France) according to the manufacturer’s instructions. The blood parameters recorded were: white blood cells, lymphocytes, monocytes, granulocytes, red blood cells and haematocrit. Samples were processed in duplicate.

A general linear model (GLM) was fitted for the analysis of BW in each week, including as fixed effect the experimental group with 2 levels (VT and NC) and the covariate number of liveborn at birth. Differences in BW in each week between the experimental groups were computed and plotted as the difference in their least squares means ± standard error of means (NC-VT). As previously, for organ weight and blood parameters analysis a GLM was fitted including as fixed effect the experimental group (VT and NC), but in the case of organ weights data were corrected using BW as covariate.

Growth curves for male rabbits were fitted following the Gompertz model as described by Laird [45], as the Gompertz curve is appropriate to describe rabbit growth [43]. The parameters of the Gompertz curve were estimated and differences between experimental groups for the Gompertz curve parameters were tested using GLM as previously. Differences of p<0.05 were considered significant. Statistical analyses were performed with SPSS 21.0 software package.

### Sampling for the molecular signature of hepatic tissue

A total of 6 samples (3 VT and 3 NC) were generated obtaining some liver biopsies randomly. The samples were immediately washed with DPBS to remove blood remnants. After that, one part from each sample was stored in RNA-later (Ambion Inc., Huntingdon, UK) at -20 °C for transcriptomic analysis, while the other part was directly flash frozen in liquid nitrogen and stored at -80° C for the proteomic study. Then, all samples were derived from the same animal cohorts.

### Transcriptome: RNA isolation, RNA-Seq and functional analysis

Samples were shipped to the Macrogen company (Seoul, South Korea). Afterwards, the mRNA purification was carried out using Sera-mag Magnetic Oligo (dT) Beads, followed by buffer fragmentation. Reverse transcription was followed by PCR amplification to prepare the samples to be sequenced, keeping the strand information, in an Illumina Hiseq-4000[D1] sequencer (Illumina, San Diego, USA). Resulting raw sequences are available at the NCBI Sequence Read Archive (BioProject ID: PRJNA483095, Supplemental Table 1). Raw read qualities were assessed using FastQC software [46]. Reads were mapped keeping strandness information against the reference genome for *Oryctolagus cuniculus*, version 2.0 from Ensembl using hisat2 [47]. Expression was counted using stringtie [48]. This counting was guided using the genome annotation and a unified set of transcripts was created for the samples analysed. Then, a Fragments Per Kilobase of transcript per Million (FPKM) table with gene expression for each sample was generated and used for assess the expression profiles of each sample by PCA. Then, a table with raw counts were generated. This table was used for the differential expression analyses using edgeR [49] integrated in the webservice platform WebMeV [50]. Only differential expressed genes (DEG) with a threshold of a false discovery rate (FDR) of < 0.05 were considered for further analyses.

**Table 1.**
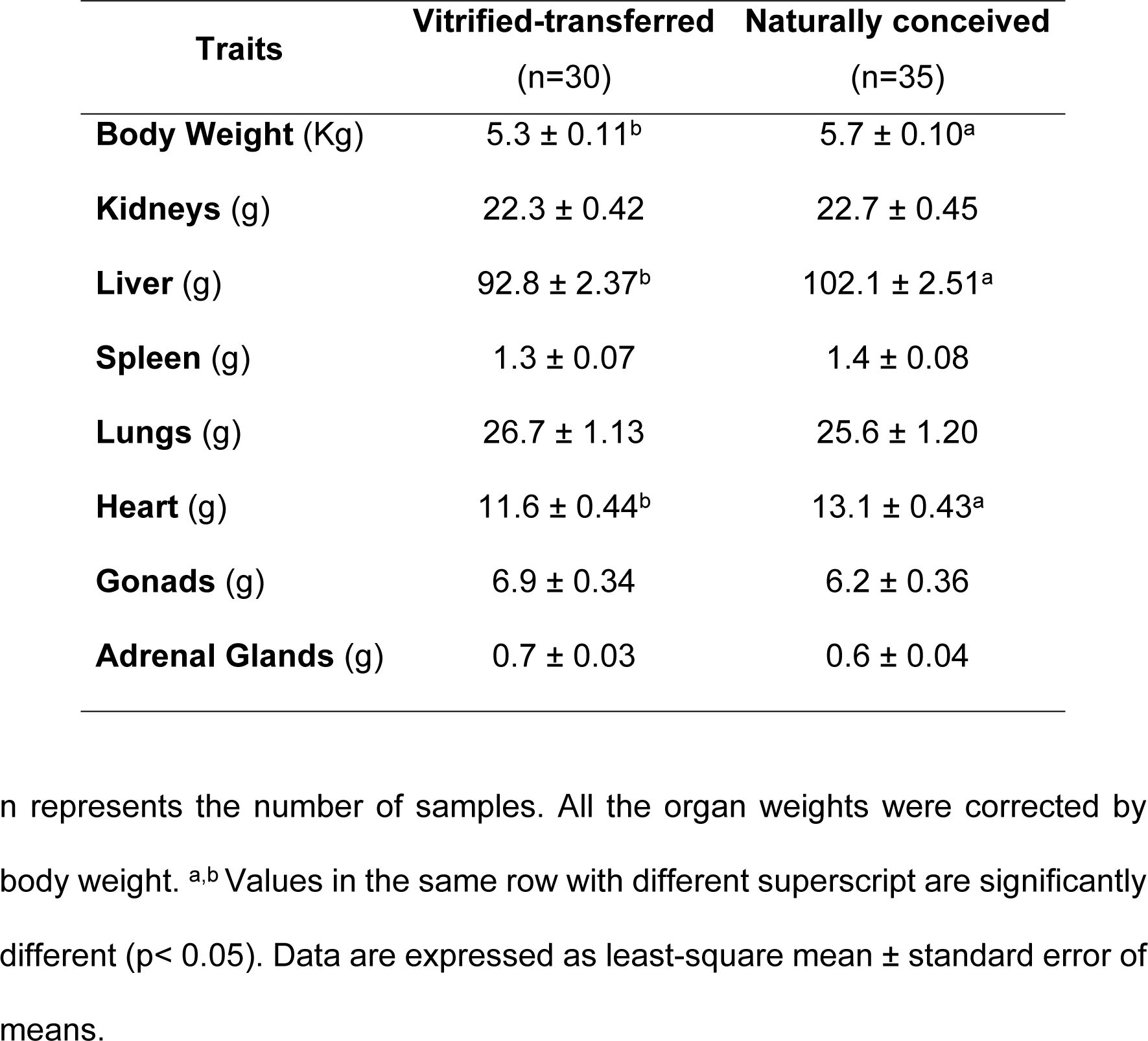
Body weight and dissection data of adult males derived from vitrified-transferred embryos and naturally conceiving.

For comparison between groups, further filtering of DEGs was made. Only genes with an absolute value of log2 fold change => 1.2 and at least one of the samples involved in the comparison with a Transcripts Per Kilobase Million (TPM) of 1 were kept. Then, in those samples which registered a coefficient of variation higher than 50% and a difference between mean and median higher than 1, the gene was maintained if half of the samples of the most expressed condition group had an expression two times higher than the mean of the other group.

ClustVis software was used to perform the Principal Component Analysis (PCA) of all expression data and the Heat-Map clustering [51]. Functional annotation of DETs, enrichment analysis of their associated GO terms and KEGG pathways analysis were computed using the Bioinformatic software: David Functional Annotation Tool (version 6.8; October 2016), considering a P-value < 0.05.

### Proteome: Protein extraction, identification and functional analysis

Here we maintained the same cohort of animals used for the transcriptomic study. Samples were placed in 8M urea (Malinckrodt AR^®^, LabGuard^®^) in homogenisation tubes (^RT^Precellys^®^ Ceramic Bead Tube) and then ground using the homogeniser PrecellysTM Control Device (Bertin Technologies). Tissue extracts were subjected to cold acetone precipitation and pellets were resuspended in 8M urea, determining the protein concentration by the BCA assay kit (ThermoScientific, Meridian Rd., Rockford, IL, USA). A volume of sample with a protein amount of 50 µg was processed. Briefly, samples were subjected to denaturation, reduction and alkylation prior to the digestion step with trypsin/Lysine-C enzyme mix (Trypsin/Lys-C mix mass spec grade, Promega). Digested peptides were purified using C18 columns (MicroSpin Column 96/pk, C18 Silica, 5-200 µL loading, 5-60 µg capacity, The Nest Group, Inc.) and samples were dried and stored at -20 °C. Then, the resulting pellets were analysed using a Dionex UltiMate 3000 RSLC Nano System coupled to the Q Exactive(tm) HF Hybrid Quadrupole-Orbitrap Mass Spectrometer (Thermo Scientific, Waltham, MA, USA) as described in [52].

The peptide masses were searched against a protein database for the taxa *Oryctolagus cuniculus* (UniProt) using the freely available MaxQuant software package (version 1.5.5.1, Max Planck Institute of Biochemistry), with first search peptide tolerance of 20 ppm, main search peptide tolerance of 4.5 ppm, 1% false discovery rate (FDR), trypsin and lysC digestion and carbamidomethyl cysteine as fixed modification, and oxidised methionine as variable modification. Match between runs was considered (Match time window of 1 min and an Alignment Time Window of 20 min). Label-free quantification (LFQ) was used to obtain the normalised LFQ intensity. Contaminants and reverse proteins were removed from the analysis. Only proteins with at least 2 MS/MS counts were considered.

InfernoRDN application (Pacific Northwest National Laboratory), which provides an easy-to-use R (version 3.3.1) for proteomic data analysis, was used to analyse and compare the intensity and the LFQ intensity data, including Log2 transformation and Analysis of Variance (ANOVA). Only proteins that displayed values of intensity≠0 in the 2 control samples and at least 3 vitrified samples were included in the ANOVA. After the InfernoRDN analysis, proteins with ANOVA p-value<0.05 were subjected to cross comparison between groups (control vs vitrified). Proteins with an average fold change (FC)≥2 or ≤0.5, or with a t-test<0.05 (and a FC ≥1.5 or ≤0.66), were selected for the subsequent functional annotation.

ClustVis software was used to perform the Principal Component Analysis (PCA) of all expression data and the Heat-Map clustering [51]. Functional annotation of differentially expressed proteins (DEPs), enrichment analysis of their associated GO terms and the KEGG pathways analysis were computed using the Bioinformatic software: David Functional Annotation Tool (version 6.8; October 2016), considering a P-value < 0.05.

## Results

### Males derived after vitrified embryo transfer procedure exhibit lower growth performance and lower body and organ weights in adulthood

Although at parturition the VT group showed higher individual weight than NC group (67.8±1.46 vs 60.5±1.72 g, p<0.05), even after data were corrected by number of born offspring (6.9±0.38, significant covariate effect at p<0.05), a reduced growth was observed from the second week of life until adulthood was reached (Figure 2 and Figure 3). Hence, the parameters governing the Gompertz growth curve (Figure 2) revealed that its estimated values; related with the initial body weight condition (b parameter: 4.55 ± 0.124 vs 5.23 ± 0.209, p<0.05), growth velocity (k parameter: 0.16 ± 0.005 vs 0.20 ± 0.007, p<0.05) and mature body weight (a parameter, 4873.2 ± 82.32 vs 5275.5 ± 105.20, p<0.05) were lower in the VT vs NC group, respectively. After weaning (from 4 to 9 weeks of age), mean weight differences between groups were 248.0±20.98 g (NC-VT ± standard error, p<0.05, Figure 3). From this age, the weight differences were still increased until adulthood (from 10 to 56 weeks of age), the mean weight differences between groups being 696.8±44.42 g (NC-VT ± standard error, p<0.05, Figure 3).

**Fig 2.**
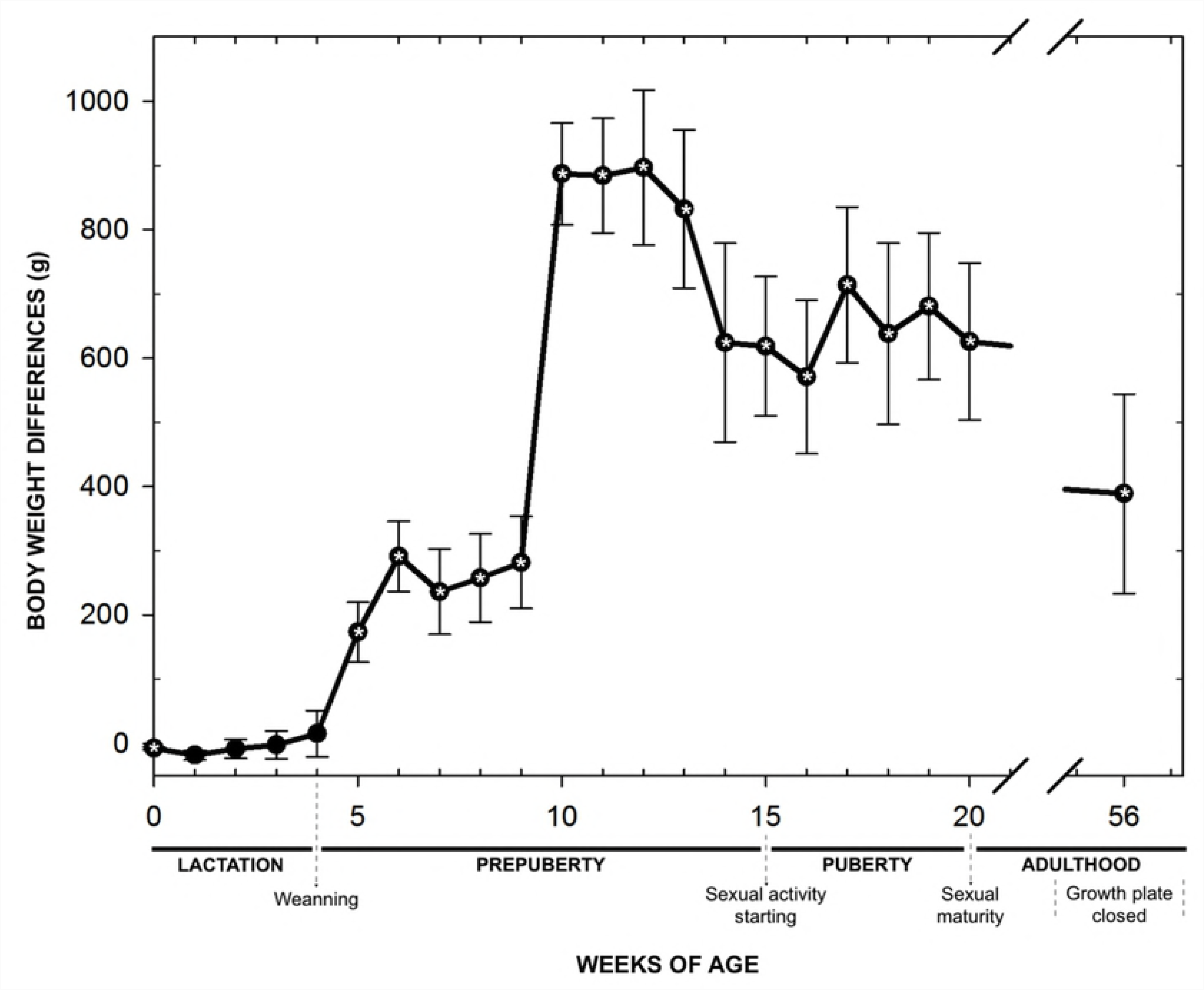
Growth curves from vitrified-transferred embryos and naturally conceiving.

**Fig 3.**
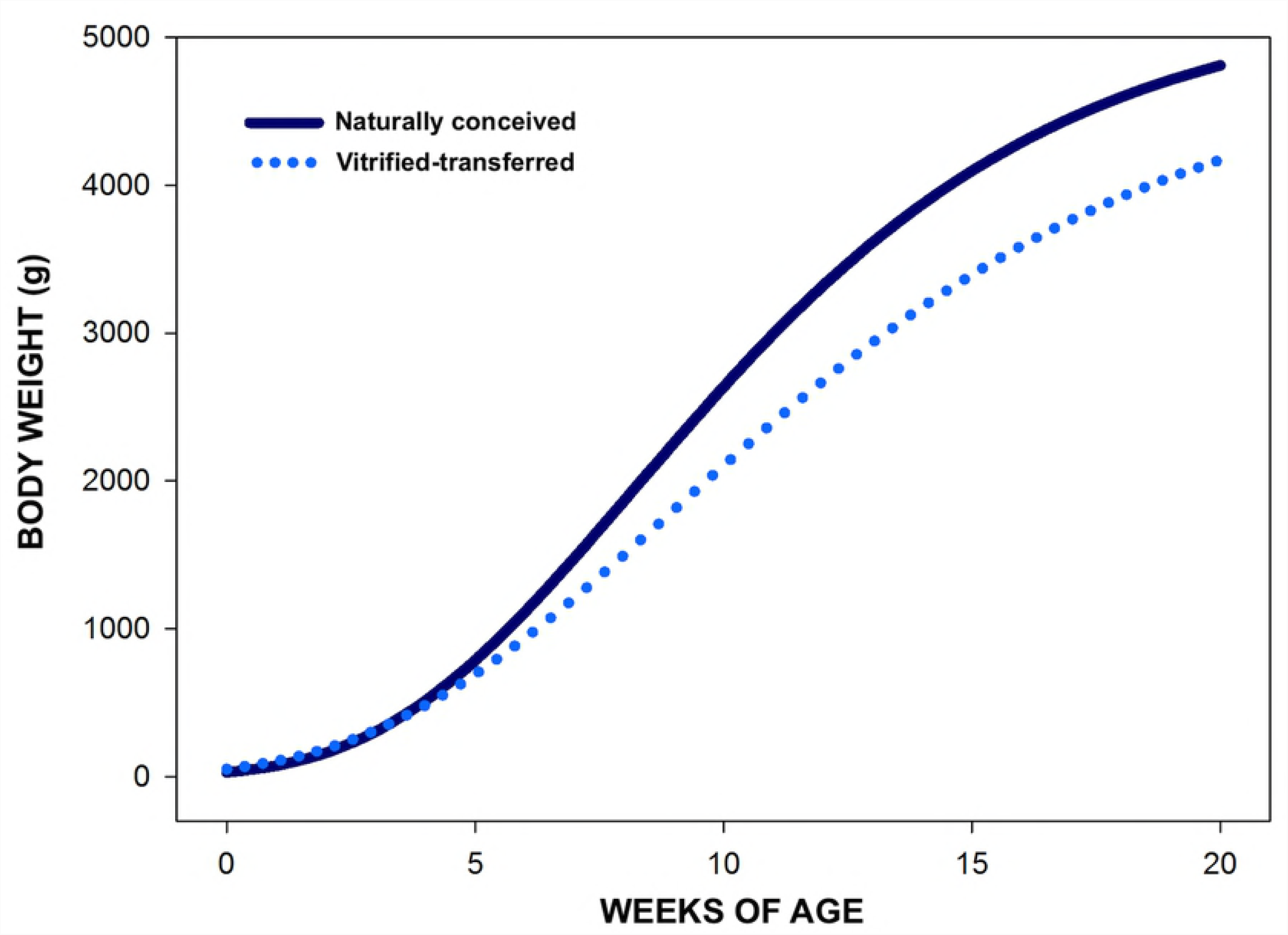
Differences in body weight between naturally conceived (NC) and vitrified-transferred (VT) groups during development, computed as NC-VT.

At 56 weeks of age, VT group showed lower body weight (5.3 ± 0.11 vs 5.7 ± 0.10 Kg, for VT vs NC group respectively, p<0.05, Table 1). Moreover, VT group showed lower liver (92.8 ± 2.37 vs 102.1 ± 2.51 g [10.0%], for VT vs NC group respectively, p<0.05) and heart weight (11.6 ± 0.44 vs 13.1 ± 0.43 g [12.9%], for VT vs NC group respectively, p<0.05), even after data were corrected by body weight (Table 1). No significant weight differences were observed for the rest of the organs analysed.

### Vitrified embryo transfer procedure seems to be neutral on the peripheral blood parameters

As shown in Table 2, there were no significant differences in peripheral profile of blood cells (white blood cells, red blood cells, and haematocrit), between VT and NC groups.

**Table 2.** Haematological comparison between vitrified-transferred and naturally conceived groups.

| Parameters | Vitrified-transferred<br>(n=10) | Naturally conceived<br>(n=10) | p-value |
| --- | --- | --- | --- |
| **WBC** ($10^3/\text{mm}^3$) | 12.2 ± 1.49 | 13.6 ± 1.49 | 0.491 |
| **LYM** ($10^3/\text{mm}^3$) | 6.8 ± 1.46 | 7.7 ± 1.46 | 0.663 |
| **MON** ($10^3/\text{mm}^3$) | 1.6 ± 0.143 | 1.8 ± 0.143 | 0.221 |
| **GRA** ($10^3/\text{mm}^3$) | 3.8 ± 0.38 | 4.15 ± 0.38 | 0.573 |
| **RBC** ($10^6/\text{mm}^3$) | 5.8 ± 0.095 | 5.9 ± 0.095 | 0.866 |
| **HTO** (%) | 40.6 ± 1.17 | 41.1 ± 1.17 | 0.812 |

WBC: White Blood Cells; LYM: Lymphocytes, MON: Monocytes, GRA: Granulocytes ( $10^3/\text{mm}^3$ ), RBC: Red Blood Cells, HTO: Haematocrit. Data are expressed as least-square means ± standard error of means.

### The liver transcriptome was influenced by vitrified embryo transfer procedure

The transcriptomes from adulthood liver tissue from VT were compared with NC counterparts. The mean number of raw reads was 48.62 ± 4.48 (±SD) millions, and transcripts from 13.908 to 14.524 different genes (from a total of 24.964 annotated transcripts of Orycun2.0) were detected in each individual. Principal Component Analysis (PCA) and Heat-Map analysis showed that, while the NC samples showed higher variability, the VT samples clustered together (Figure 4A, 4B). RNA-Seq data analysis identified 133 differentially expressed transcripts (DETs) between the VT and NC groups. Of the transcripts that were significantly different, a total of 96 DETs were recognised by the DAVID bioinformatics tool. From these DETs, there was a higher number of downregulated (68/96, [70.8%]) than up-regulated (28/96, [29.2%]) in VT samples compared with NC group. A description of DETs and the fold change values obtained are shown in Table 3. Functional GO term enrichment and KEGG pathway analysis of DETs were recorded in Table 4. This analysis suggests transcriptomic alteration related to zinc homeostasis, lipid metabolism and hepatic immune pathways in VT group animals.

**Table 3.**
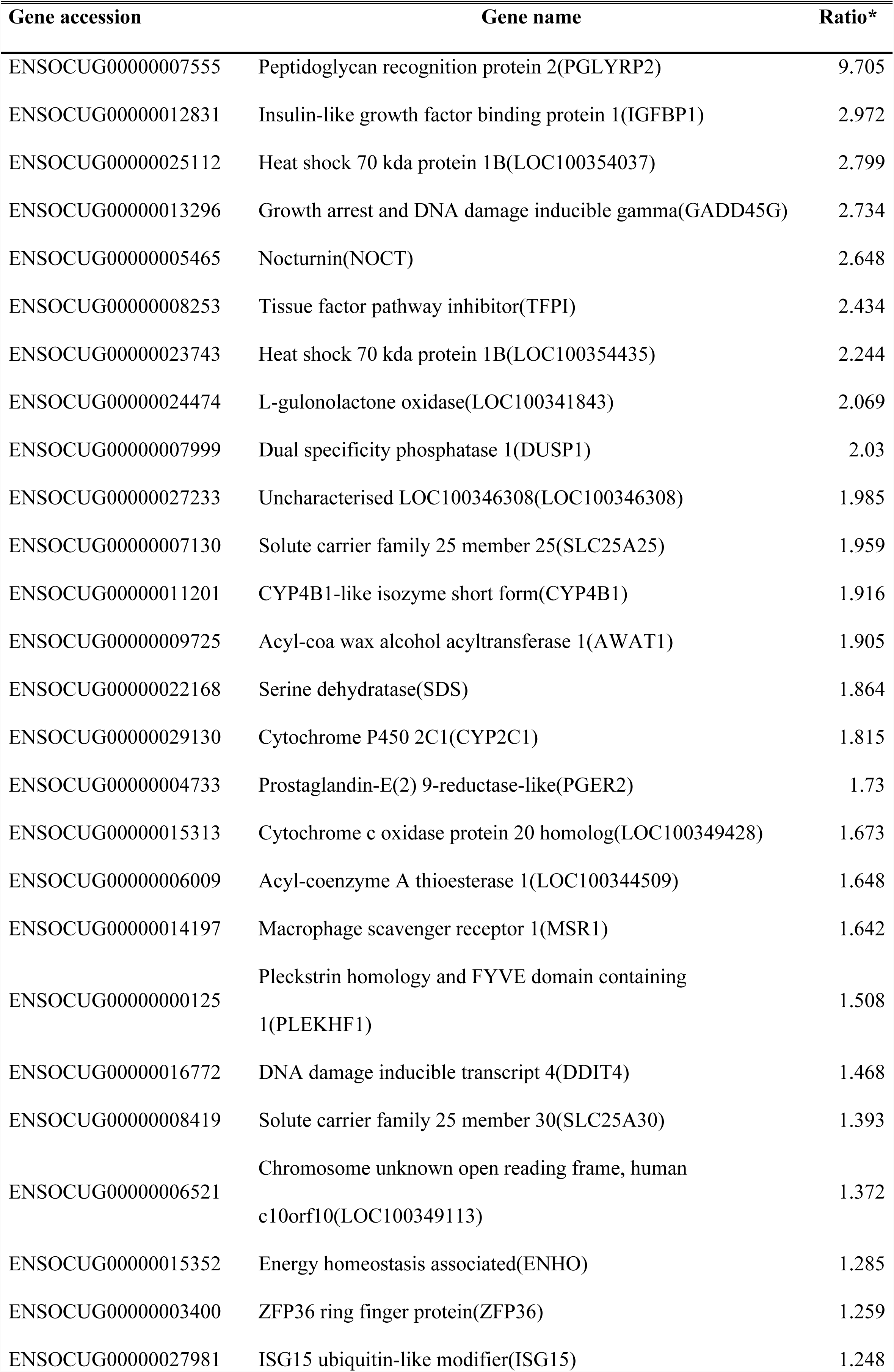
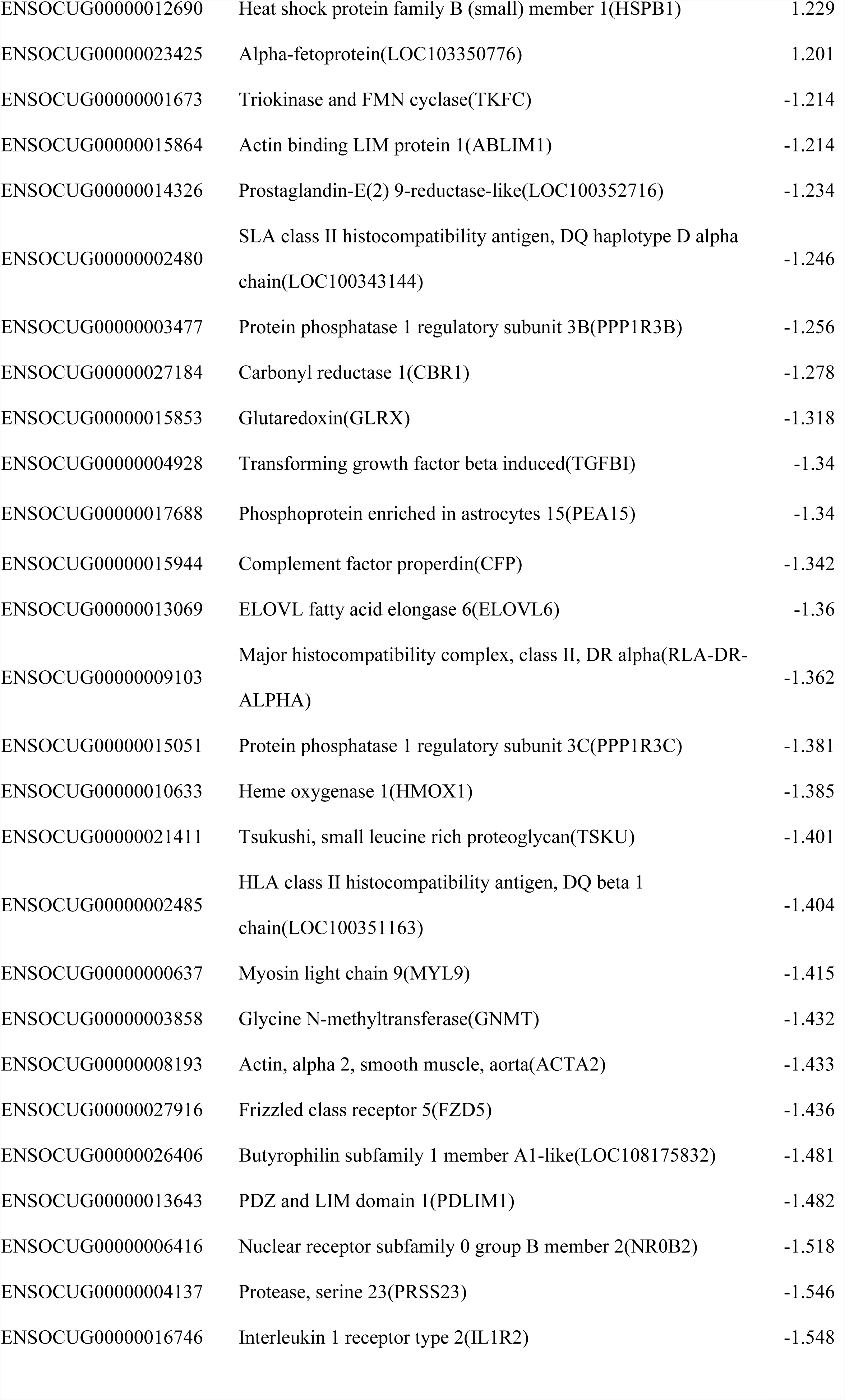
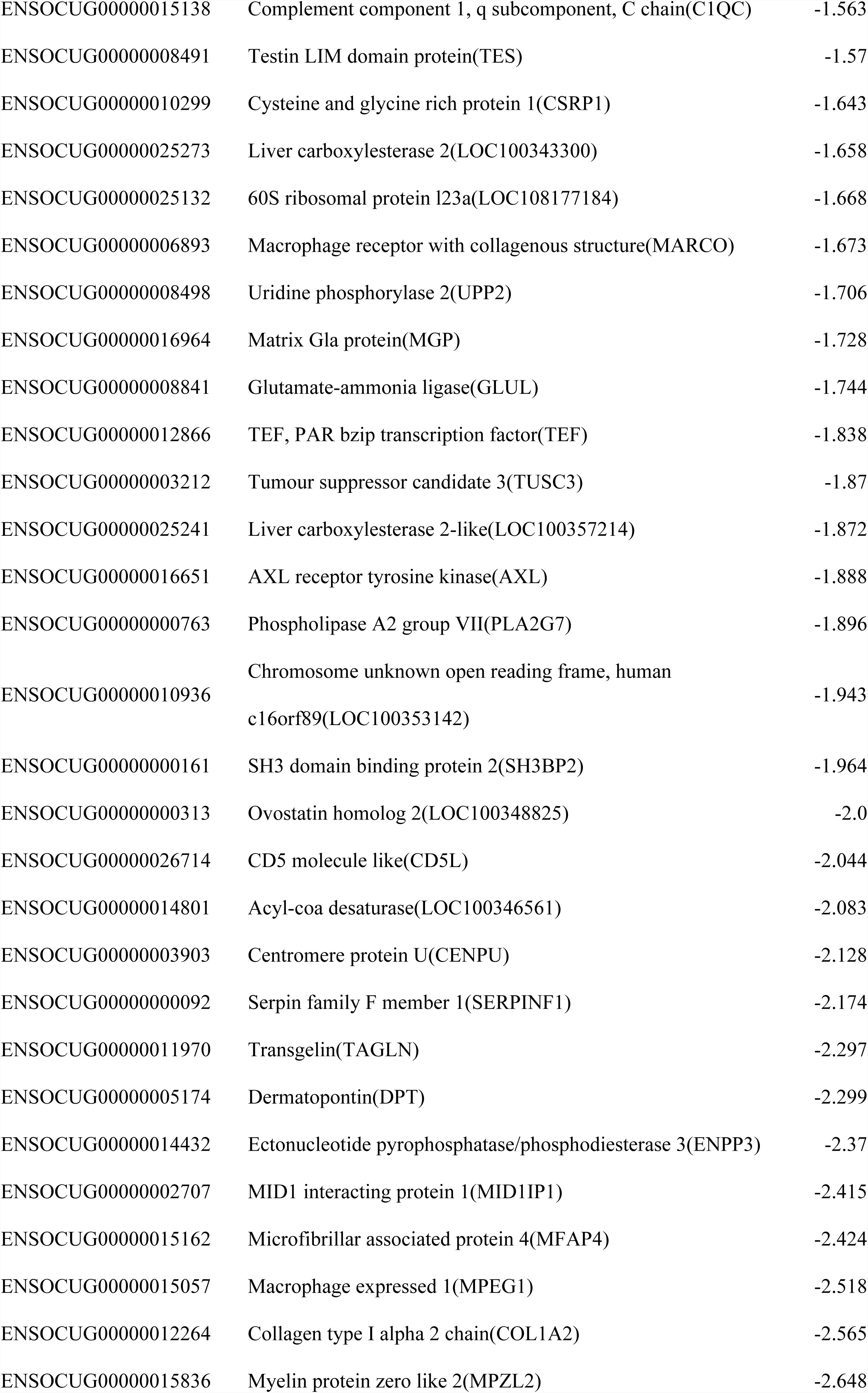
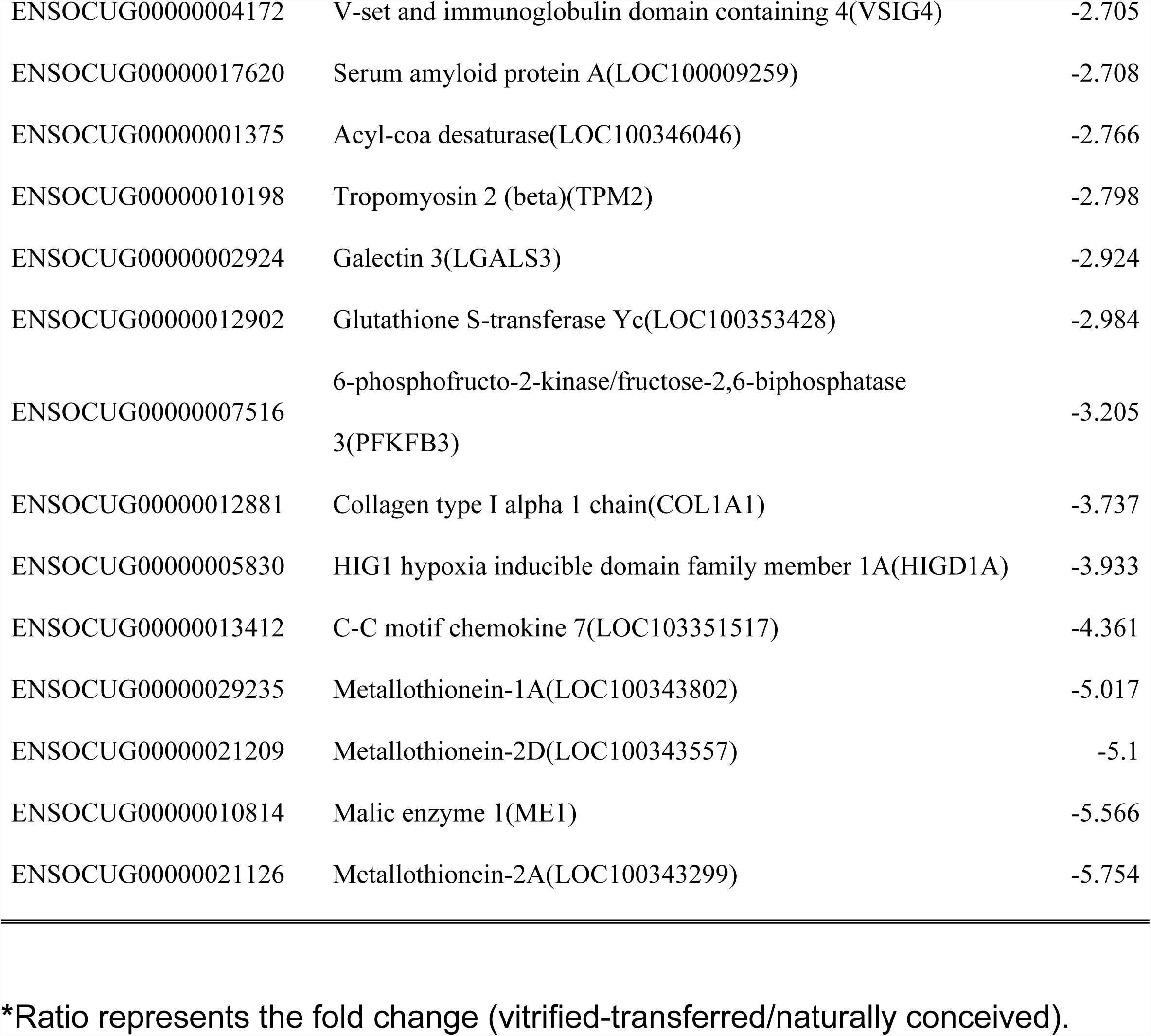
Differentially expressed transcripts in the liver tissue.

**Table 4.**
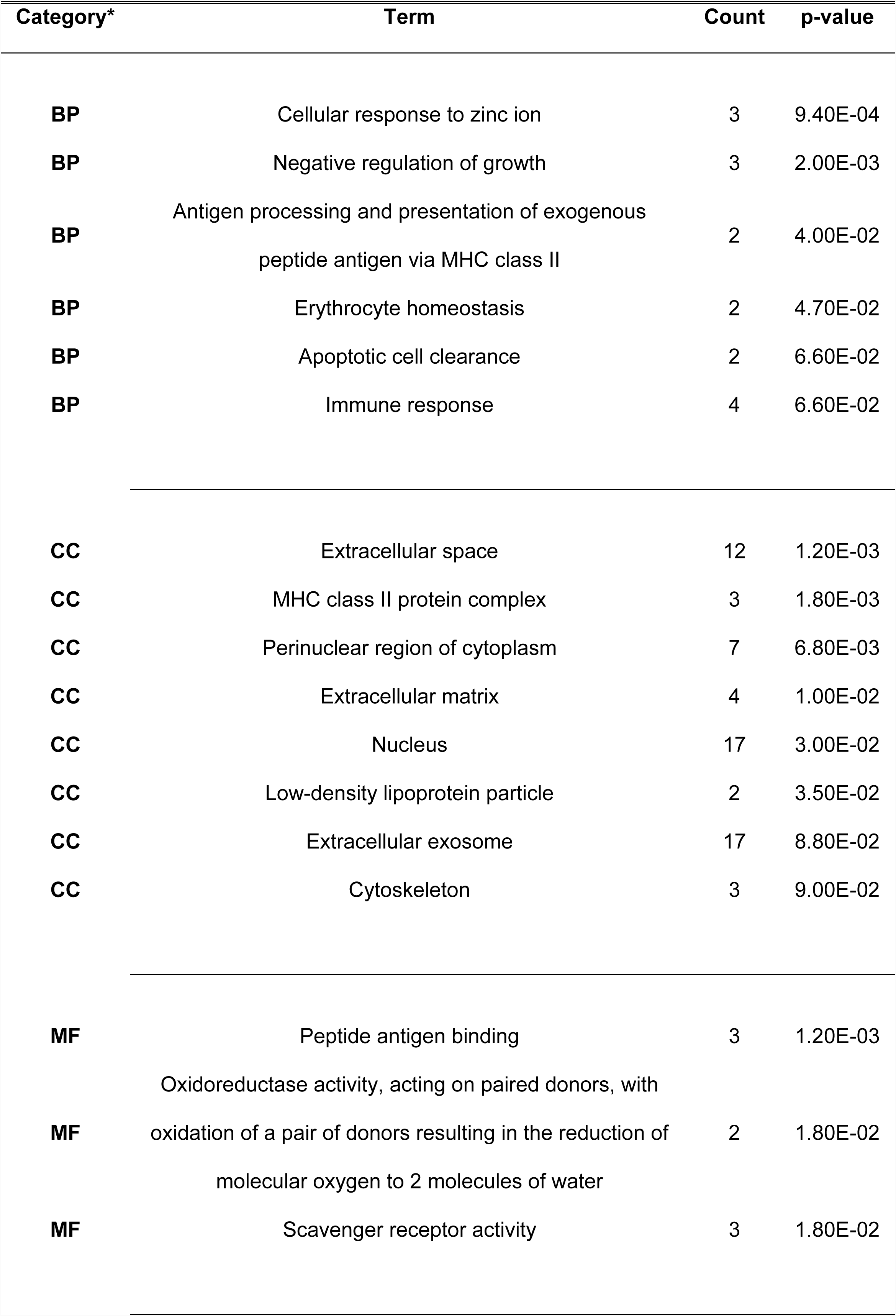
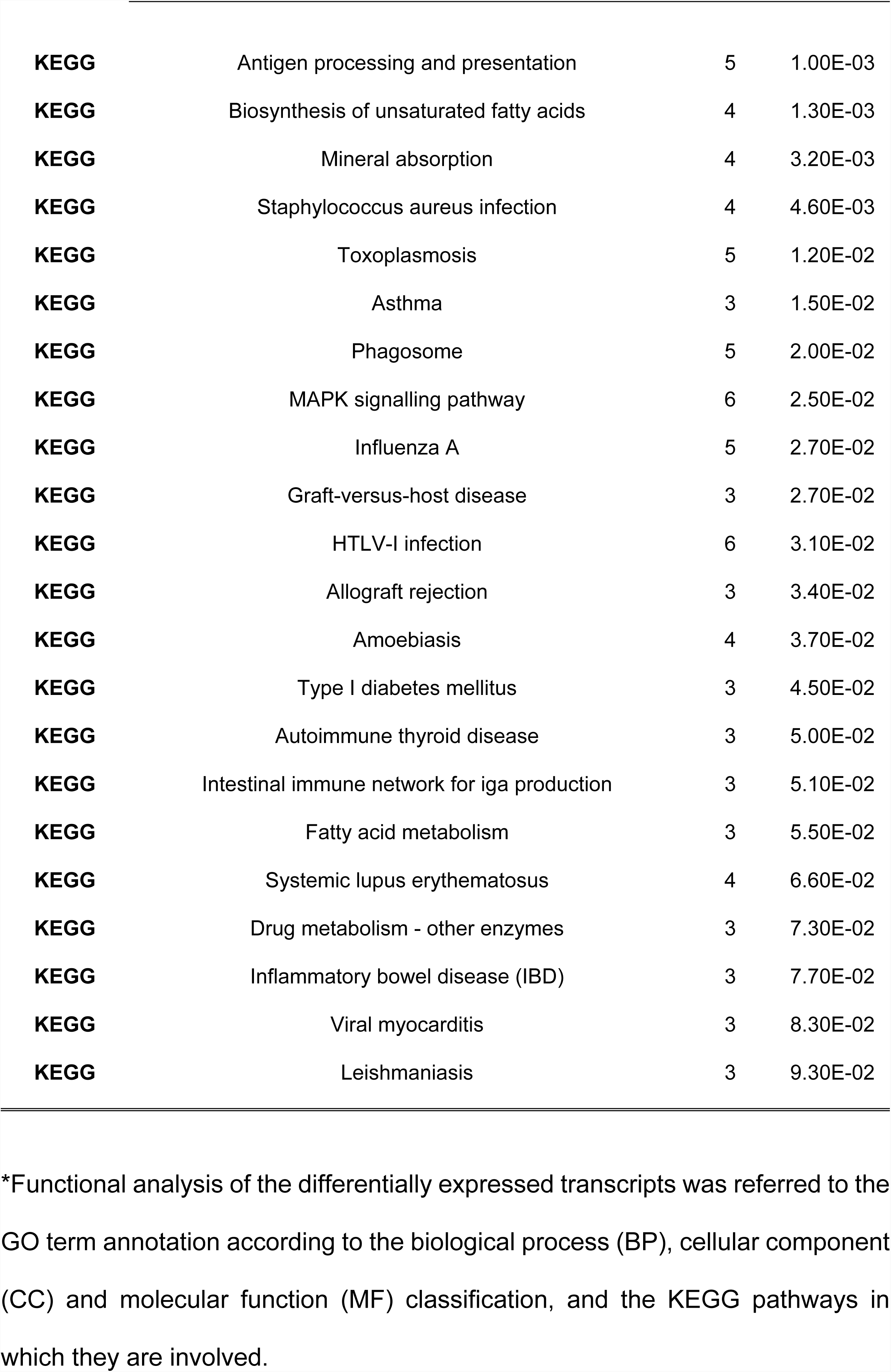
Functional analysis of the differentially expressed transcripts.

| Category* | Term | Count | p-value |
| --- | --- | --- | --- |
| BP | Cellular response to zinc ion | 3 | 9.40E-04 |
| BP | Negative regulation of growth | 3 | 2.00E-03 |
| BP | Antigen processing and presentation of exogenous peptide antigen via MHC class II | 2 | 4.00E-02 |
| BP | Erythrocyte homeostasis | 2 | 4.70E-02 |
| BP | Apoptotic cell clearance | 2 | 6.60E-02 |
| BP | Immune response | 4 | 6.60E-02 |
| CC | Extracellular space | 12 | 1.20E-03 |
| CC | MHC class II protein complex | 3 | 1.80E-03 |
| CC | Perinuclear region of cytoplasm | 7 | 6.80E-03 |
| CC | Extracellular matrix | 4 | 1.00E-02 |
| CC | Nucleus | 17 | 3.00E-02 |
| CC | Low-density lipoprotein particle | 2 | 3.50E-02 |
| CC | Extracellular exosome | 17 | 8.80E-02 |
| CC | Cytoskeleton | 3 | 9.00E-02 |
| MF | Peptide antigen binding | 3 | 1.20E-03 |
| MF | Oxidoreductase activity, acting on paired donors, with oxidation of a pair of donors resulting in the reduction of molecular oxygen to 2 molecules of water | 2 | 1.80E-02 |
| MF | Scavenger receptor activity | 3 | 1.80E-02 |
| **KEGG** | Antigen processing and presentation | 5 | 1.00E-03 |
| **KEGG** | Biosynthesis of unsaturated fatty acids | 4 | 1.30E-03 |
| **KEGG** | Mineral absorption | 4 | 3.20E-03 |
| **KEGG** | Staphylococcus aureus infection | 4 | 4.60E-03 |
| **KEGG** | Toxoplasmosis | 5 | 1.20E-02 |
| **KEGG** | Asthma | 3 | 1.50E-02 |
| **KEGG** | Phagosome | 5 | 2.00E-02 |
| **KEGG** | MAPK signalling pathway | 6 | 2.50E-02 |
| **KEGG** | Influenza A | 5 | 2.70E-02 |
| **KEGG** | Graft-versus-host disease | 3 | 2.70E-02 |
| **KEGG** | HTLV-I infection | 6 | 3.10E-02 |
| **KEGG** | Allograft rejection | 3 | 3.40E-02 |
| **KEGG** | Amoebiasis | 4 | 3.70E-02 |
| **KEGG** | Type I diabetes mellitus | 3 | 4.50E-02 |
| **KEGG** | Autoimmune thyroid disease | 3 | 5.00E-02 |
| **KEGG** | Intestinal immune network for iga production | 3 | 5.10E-02 |
| **KEGG** | Fatty acid metabolism | 3 | 5.50E-02 |
| **KEGG** | Systemic lupus erythematosus | 4 | 6.60E-02 |
| **KEGG** | Drug metabolism - other enzymes | 3 | 7.30E-02 |
| **KEGG** | Inflammatory bowel disease (IBD) | 3 | 7.70E-02 |
| **KEGG** | Viral myocarditis | 3 | 8.30E-02 |
| **KEGG** | Leishmaniasis | 3 | 9.30E-02 |

\*Functional analysis of the differentially expressed transcripts was referred to the GO term annotation according to the biological process (BP), cellular component (CC) and molecular function (MF) classification, and the KEGG pathways in which they are involved.

**Fig 4.**
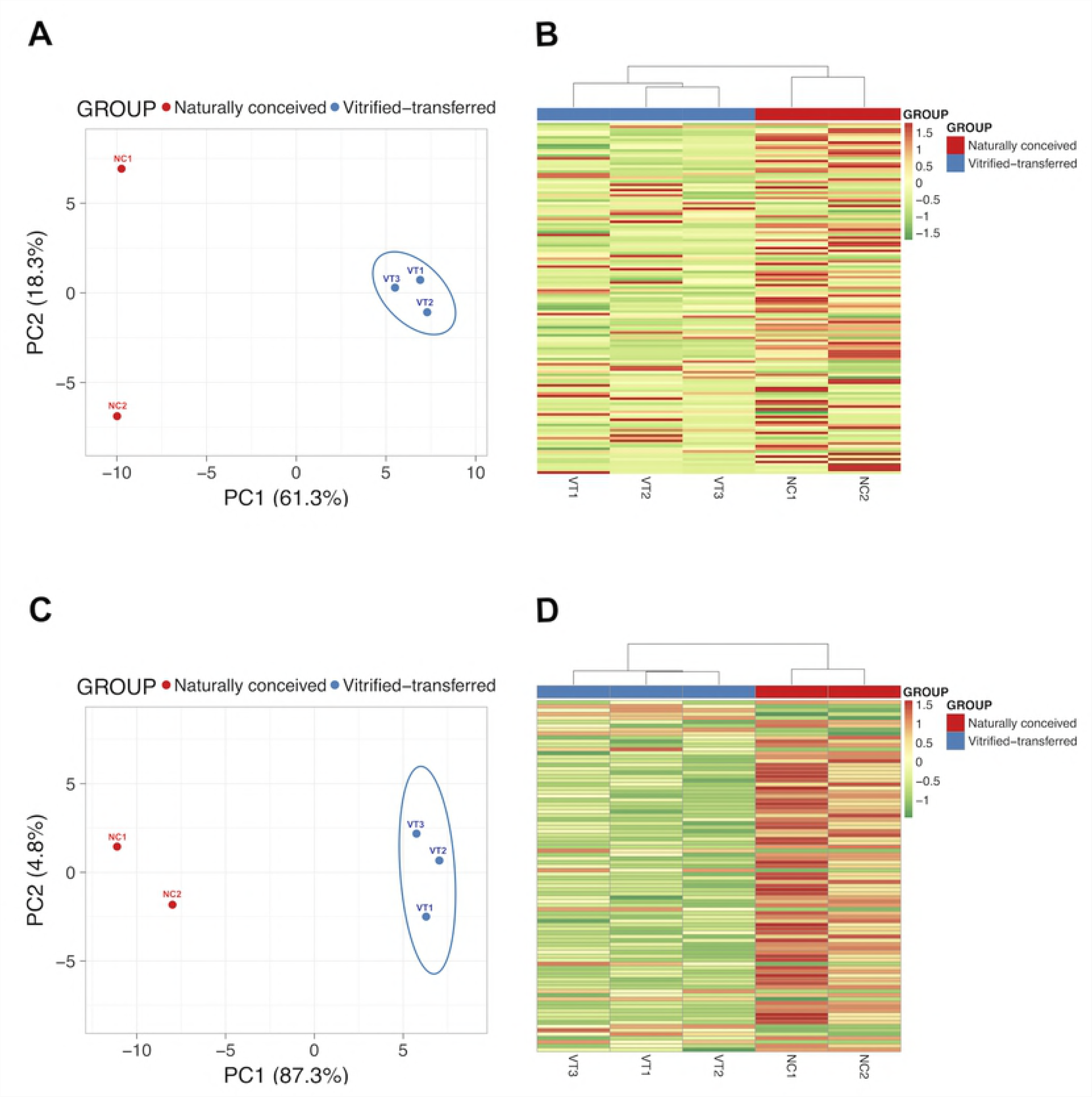
Molecular analysis in liver samples obtained from adult males derived from vitrified-transferred embryos and naturally conceiving. (A) Principal component analysis of the transcriptome. (B) Heat-Map clustering of the transcriptome. (C) Principal component analysis of the proteome. (D) Heat-Map clustering of the proteome. The representation of sample variability between the experimental groups was performed taking into account only the differentially expressed transcripts and proteins and the transcripts or proteins absent in specific groups.

### The liver protein profile was influenced by vitrified embryo transfer procedure

In this case, the protein profiles of adulthood liver tissue from VT were compared with NC counterparts. Mean number of MS/MS spectra per sample submitted for the MaxQuant analysis was 76408 ± 950 (±SD). The number of peptides identified ranged from 10708 to 12059 and the number of proteins from 1707 to 1782 (from a total of 22929 proteins included in Oryctolagus cuniculus database [Uniprot]) was detected in each individual. PCA and Heat-Map analysis showed that, despite expected individual variability, samples from each group were clustered together (Figure 4C, 4D). Protein data analysis identified 90 DEPs in VT animals compared with NC group. Of the proteins that were significantly different, a total of 76 DEPs were recognised by the DAVID software. From these DEPs, there was a higher number of downregulated (60/76, [78.9%]) than up-regulated (16/76, [21.1%]). Annotation of DEPs and the fold change values obtained are shown in Table 5. Functional GO term enrichment and KEGG pathway analysis of DEPs were recorded in Table 6. This analysis suggests an impaired oxidative metabolism related to ATP synthesis in the mitochondria in VT group animals.

**Table 5.**
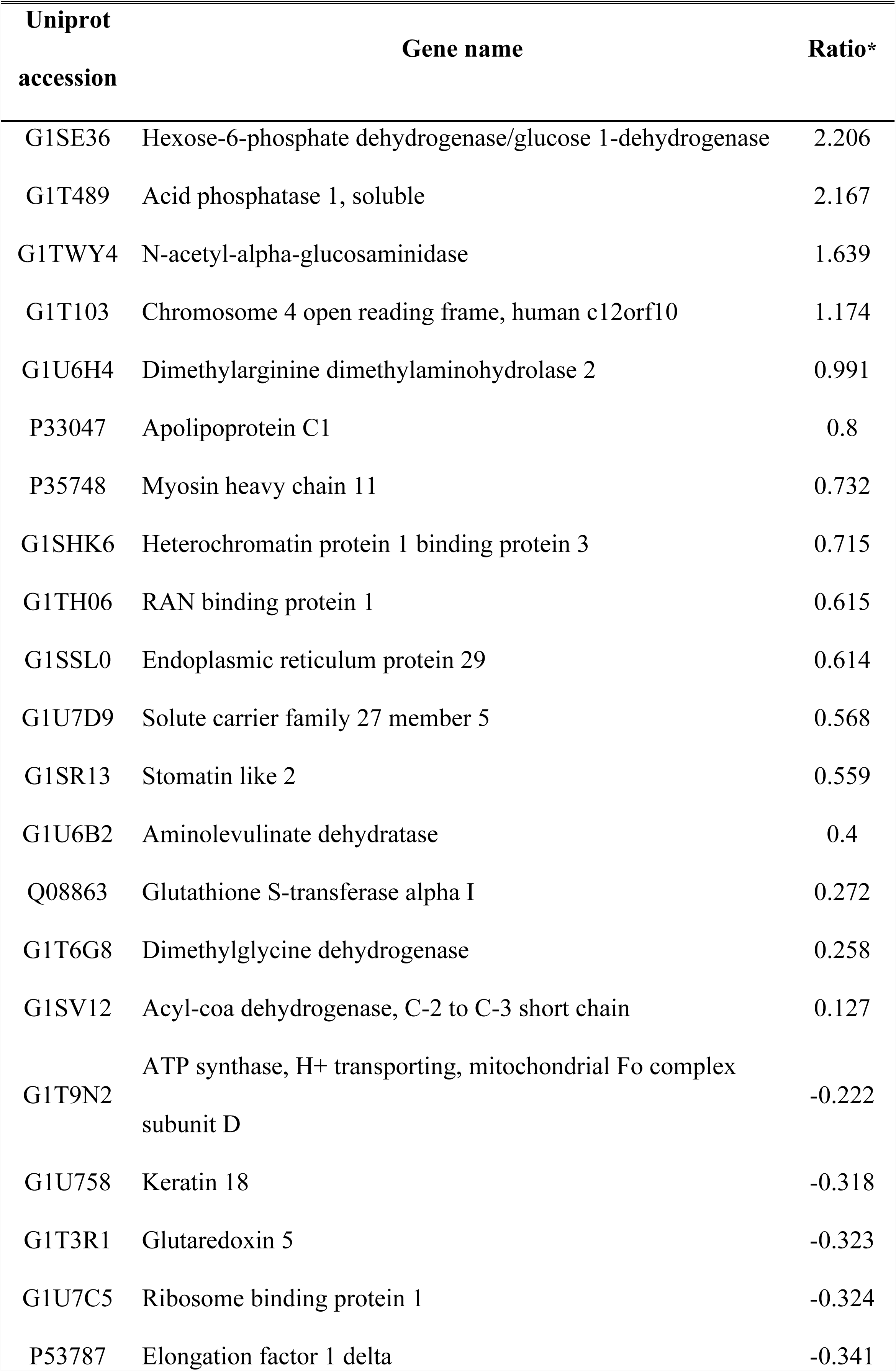
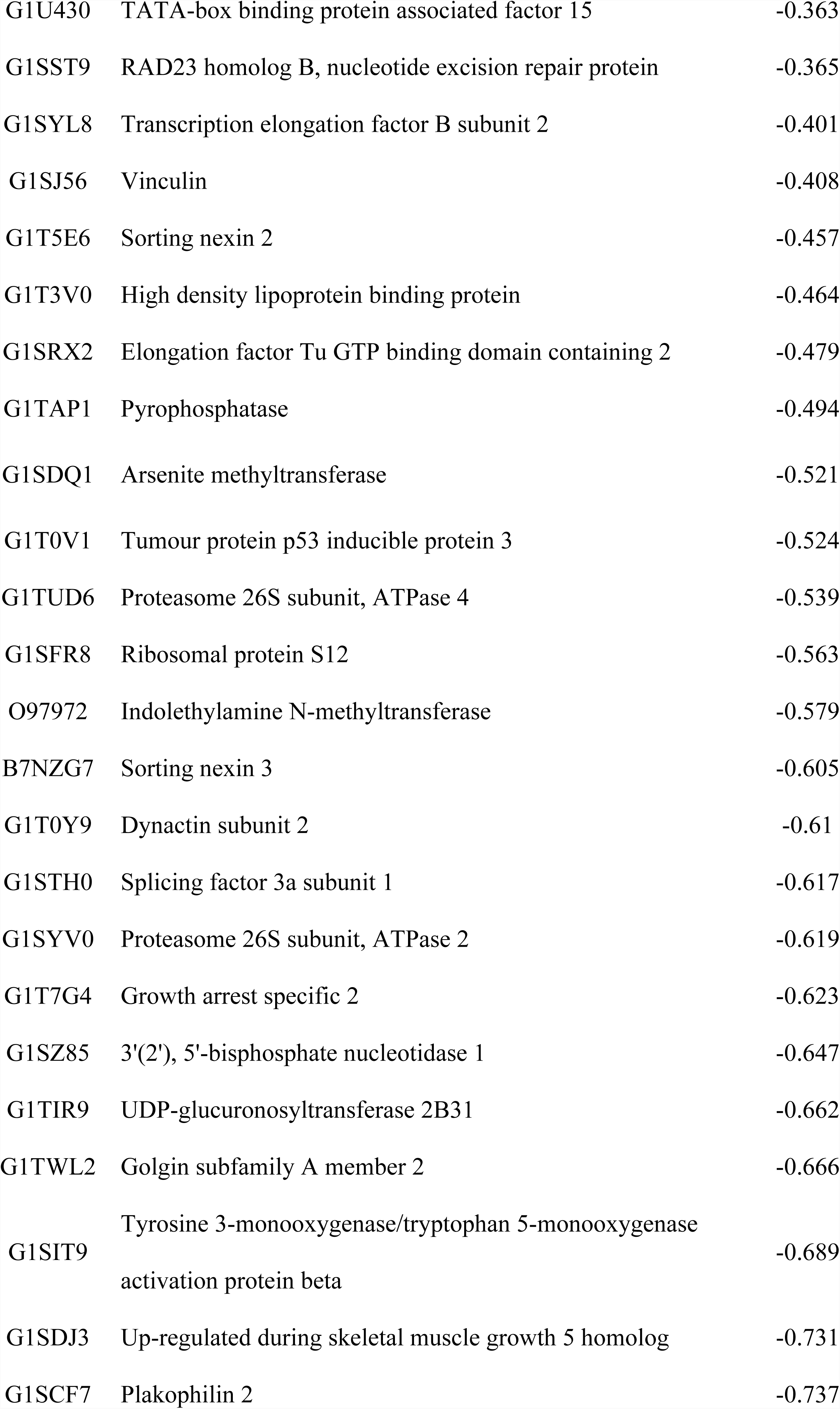
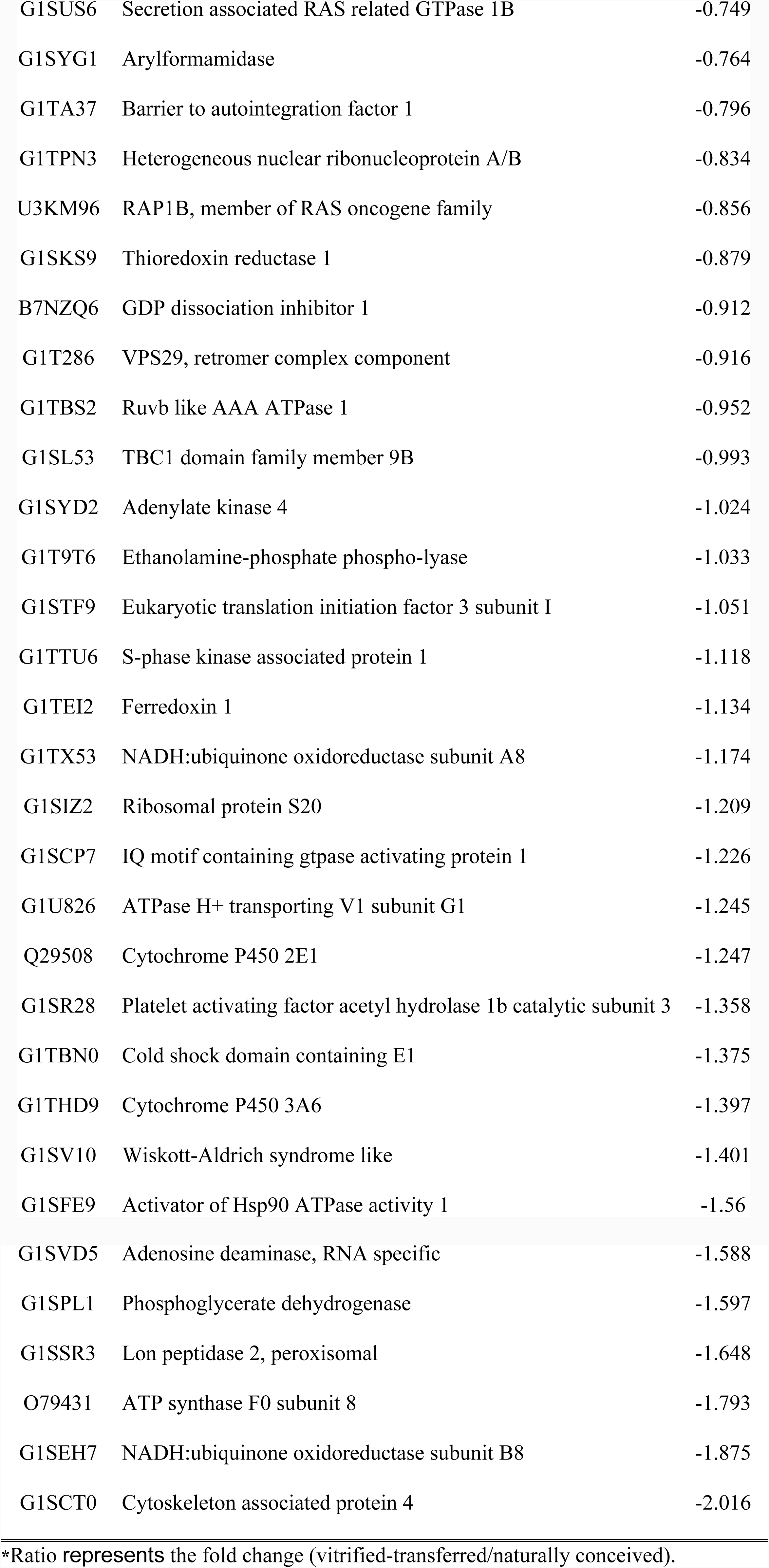
Differentially expressed proteins in liver tissue.

**Table 6.**
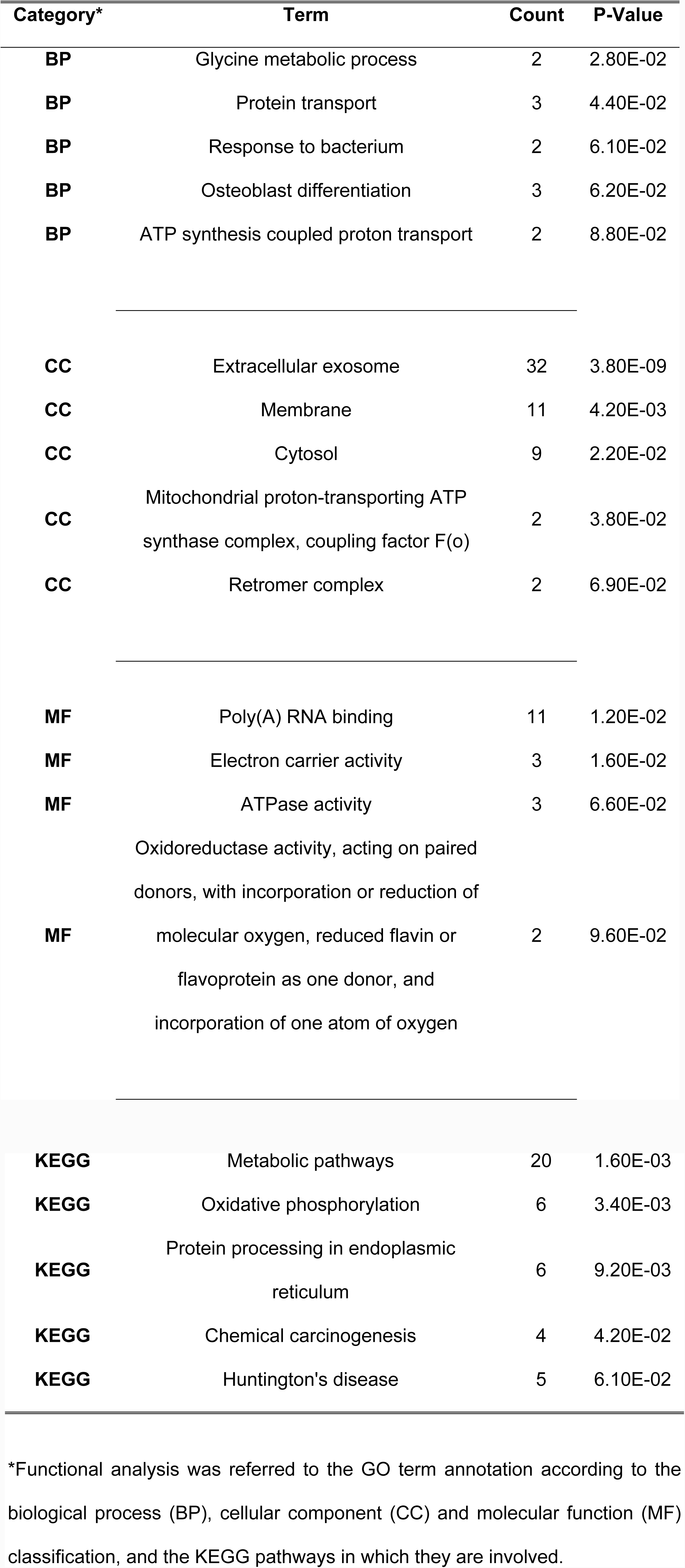
Functional analysis of differential expressed proteins.

| Category* | Term | Count | P-Value |
| --- | --- | --- | --- |
| BP | Glycine metabolic process | 2 | 2.80E-02 |
| BP | Protein transport | 3 | 4.40E-02 |
| BP | Response to bacterium | 2 | 6.10E-02 |
| BP | Osteoblast differentiation | 3 | 6.20E-02 |
| BP | ATP synthesis coupled proton transport | 2 | 8.80E-02 |
| CC | Extracellular exosome | 32 | 3.80E-09 |
| CC | Membrane | 11 | 4.20E-03 |
| CC | Cytosol | 9 | 2.20E-02 |
| CC | Mitochondrial proton-transporting ATP synthase complex, coupling factor F(o) | 2 | 3.80E-02 |
| CC | Retromer complex | 2 | 6.90E-02 |
| MF | Poly(A) RNA binding | 11 | 1.20E-02 |
| MF | Electron carrier activity | 3 | 1.60E-02 |
| MF | ATPase activity | 3 | 6.60E-02 |
| MF | Oxidoreductase activity, acting on paired donors, with incorporation or reduction of molecular oxygen, reduced flavin or flavoprotein as one donor, and incorporation of one atom of oxygen | 2 | 9.60E-02 |
| **KEGG** | Metabolic pathways | 20 | 1.60E-03 |
| **KEGG** | Oxidative phosphorylation | 6 | 3.40E-03 |
| **KEGG** | Protein processing in endoplasmic<br>reticulum | 6 | 9.20E-03 |
| **KEGG** | Chemical carcinogenesis | 4 | 4.20E-02 |
| **KEGG** | Huntington's disease | 5 | 6.10E-02 |

\*Functional analysis was referred to the GO term annotation according to the biological process (BP), cellular component (CC) and molecular function (MF) classification, and the KEGG pathways in which they are involved.

## Discussion

The progress made by ART during the past 2 decades makes a future without their use inconceivable [53]. However, it is well established that the use of these technologies has consequences on development and modification of the embryo epigenome [7, 53-56]. Hence, and in line with our previous studies, we again corroborate that male progeny born after transfer of vitrified embryos result in a reduced growth rate and vital organs weights in adulthood [34, 57]. In addition, multi-omics analyses of hepatic tissue revealed modifications in lipid metabolism and energy metabolism that could be implicated in growth and body weight. Importantly, in our study, there is strong evidence that vitrified embryo transfer manipulation technique represent a clear example of the active phenotypic plasticity exhibited by the embryo, where irreversible phenotypic variation in traits of individuals induce modifications of development and growth [58].

Today, cryopreservation is an essential component in the treatment of patients undergoing ART [59, 60]. The choice of a cryopreserved cycle avoids the suboptimal endometrium generated by supra-physiologic hormonal levels during a conventional ovarian stimulation [61, 62], improving the implantation and pregnancy rates and reducing the risk of preterm birth [63, 64]. However, although cryopreservation was considered a neutral technique for years [65, 66], some recent studies have revealed potential adverse effects throughout gestation [24, 25, 67, 68]. Associations between ART and birth defects or stillbirth are controversial, but the vast majority of studies indicated that the increased risk of birth defects are attributed to maternal characteristics related to infertility [69-72]. However, in recent years, research studies are starting to learn about the long-term health of people conceived after ART treatment, but there are only few longitudinal studies about the effect of embryo cryopreservation on health risks [73-75]. To test the effect of embryo vitrification in adulthood, we used an animal model to minimise external confounding factors. It is important to state that, in our study, the offspring born after transfer of vitrified embryos were apparently healthy, which was also corroborated in adulthood by outcomes of peripheral blood features. The most remarkable finding regarding to long-term consequences of vitrification at late adulthood was lower growth, decreased body weight and a lower weight of some vital organs, such as the liver and heart. Even though higher body weight was observed at parturition, from the third-fourth week of age till adulthood animals born after transfer of vitrified embryos showed a reduced body growth curve, although in rabbit these deviations can be restored at adulthood through compensatory growth [76]. Studies on health outcomes of offspring conceived by ART in animals [77] and human [78, 79] also revealed a significant increase in the birth weight. This may be related to the fact that embryos are grown for one extra day *in vitro* after thawing to compensate for the loss of cells in the freezing and thawing processes [79]. Imprinting modifications of some growth-related genes have also been suggested to explain the phenotype variations observed in animals obtained after embryo cryopreservation [57,79]. Supporting this, previous studies have also reported variations in body weight after embryo cryopreservation [23, 57, 80], as well as after *in* vitro culture and transfer [81]. From these studies, we can learn that different ARTs protocols would lead to different outcomes via specific epigenetic modifications. Collectively, these findings suggest that features exhibited vitrified embryo transfer manipulation technique seems to be a clear manifestation of the embryonic active phenotypic plasticity, which refers to the capacity of a genotype to produce different phenotypes in response to environmental variation, contributing to diversity among individuals, populations and species [58].

Today, the ‘omics’ sciences are used to describe the flow of biological information in an organism [82]. Based on the phenotyped changes observed in adulthood of animals born after transfer of vitrified embryos, we assessed the molecular signatures at hepatic level in order to find out the possible vitrification effects on both transcriptome and proteome profile “what appears to be happening” (transcriptome) and “what makes it happen” (proteome). In our study, we registered significant differences in both levels, 133 genes and 90 proteins.

Furthermore, the PCA revealed that animals born after transfer of vitrified embryos and animals born after natural conception formed a distant cluster. The overall result suggests that vitrification could alter hepatic function, impairing the correct establishment of the “growth hormone/insulin-like growth factor type I” axis whose perturbations are responsible for many important complications, such as growth disturbance [83-85]. Among the differential transcripts, we detected 3 metallothioneins (metallothionein-1A, metallothionein-2A and metallothionein-2D) involved in multiple interconnected signalling pathways related to “negative regulation of growth” and “cellular response to zinc ion” GO terms. Metallothioneins (MT) are small molecular weight and cysteine-rich proteins that play many important biological roles, including zinc (Zn) trafficking and potential protective effects against oxidative stress and toxic metals [86-87]. Liver is the primary storage organ of Zn, is sensitive to Zn deficiency, and is the most responsive organ for antioxidative function [88]. Zn plays a key role in growth via protein synthesis and antioxidant defence, and Zn deficiency causes growth deficits [89-90]. In concordance, previous studies have also suggested that improved growth performance may result from the role of Zn as a crucial component in the systemic antioxidative and immune network [91]. Nonetheless, KEGG analysis reveals a disturbed “mineral absorption” associated with Zn due to downregulation of MT coding genes after transfer of vitrified embryos, suggesting an impaired Zn homeostasis that can incur in the lower growth exhibited by the vitrified progeny. Similar positive correlation between Zn availability and MT expression with the growth performance has been described in farm animals and children [89, 92, 93]. As regards molecular function, DETs were highlighting the “oxidoreductase activity, acting on paired donors, with oxidation of a pair of donors resulting in the reduction of molecular oxygen to 2 molecules of water” terms, which are hierarchical and associated with desaturase enzyme activity. Desaturase and elongase enzymes play a crucial role in the metabolism regulation and biosynthesis of unsaturated fatty acids [94-96]. We observed that 2 desaturases and 1 elongase were becoming downregulated in hepatic tissue in animals obtained after transfer of vitrified embryos. Interestingly, Zn also acts as a cofactor of the fatty acids desaturase enzymes, whose metabolic roles are required for optimal growth, immune response, gene expression, visual development, neurotransmission and cognition [95, 97]. So, as expected, these findings were in concordance with the perturbed “biosynthesis of unsaturated fatty acids” and “fatty acid metabolism” highlighted by the KEGG pathways analysis, which supported a disturbed lipid metabolism in hepatic tissue due to impaired Zn metabolism after embryo vitrification. A recent report has demonstrated that ART induces modifications in the lipid metabolism in foetal hepatic tissue [18], which could be maintained in later life stages [18, 98-100], and even into adult life as our results suggest.

The most significantly increased transcript in VT animals was the peptidoglycan recognition protein 2 (*PGLYRP2*), which codes a peptidoglycan-hydrolytic amidase that participates in antimicrobial immunity, hydrolysing the biologically active peptidoglycan of the bacterial cell wall into inactive fragments [101, 102]. *PGLYRP2* is constitutively expressed in the liver in the presence of bacteria and cytokines to be secreted into blood, acting as an immunity modulator [101, 102]. Thus, higher expression levels of *PGLYRP2* may suggest compromised mechanisms against microbial infections by the immune system, whose vulnerability could facilitate infection that ultimately activates *PGLYRP2*. In concordance, a majority of the terms offered by functional enrichment and GO term analysis put the spotlight on 3 downregulated genes (SLA class II histocompatibility antigen, DQ haplotype D alpha chain; HLA class II histocompatibility antigen, DQ beta 1 chain; and major histocompatibility complex, class II, DR alpha), whose function participates in the activation of the immune response via antigen binding and presenting to the T lymphocytes [103]. Therefore, underexpression of these genes can underpin a deficient capacity for antigen presentation that could suppose immunological weakness. This situation suggests a compromised immunological function in the vitrified progeny, which ultimately may enhance the *PGLYRP2* as a consequence of higher susceptibility to microbial infections due to diminished immunological sensibility. Curiously, *PGLYRP2* have a conserved Zn^2+^-binding site in the enzyme’s catalytic groove, which is crucial for the amidase activity. Thereby, the disturbed Zn metabolism after embryo vitrification could also have an impact on the functionality of this innate immunity modulator, so higher expression of *PGLYRP2* may also be an attempt to compensate for deficient activity.

Focusing on the proteomic analysis, it also revealed some disturbed metabolic pathways related with the lipid, but also glycan, metabolism. Among the biological processes related to DEPs, we can signal the “ATP synthesis coupled proton transport”, whereas we can highlight the “ATPase activity” and the “mitochondrial proton-transporting ATP synthase complex, coupling factor F0” terms, attending to the molecular function and cellular component, respectively. KEEG pathway analysis reveals that these terms are related to some downrepresented proteins (*G1SEH7, G1U826, G1TX53, G1T9N2, O79431, G1TAP1*) involved in oxidative phosphorylation (OXPHO). This metabolic process is carried out in the mitochondria by ATP synthase complex, which is composed of 2 rotary motors, the F0 and F1 subunits, whose joint performance is required for the correct function of the complex [104, 105]. So, disturbed ATP synthase activity suggests a mitochondrial dysfunction that could impair the *OXPHO* and ATP production. In agreement, Feuer *et al.* [16] observed that mitochondrial dysfunction and *OXPHO* changes were exhibited in the livers after IVF conception, reporting changes in the ATP levels. It is important to indicate that *OXPHO* plays a key role in processes such as energy production, generation of free radicals and apoptosis [106], whose disturbances are found alone or in combination in most human diseases such as intrauterine growth retardation, prematurity, low birth weight, poor weight gain, major growth retardation, short stature and dwarfism [107-110]. Intriguingly, improvements in this mitochondrial function attenuated the postnatal energy deficiency and resulted in normalisation of body weight gain [100]. Assuming this, DEPs involved in *OXPHO* suggested a disturbed energy metabolism that might explain the lower growth curve presented by rabbits obtained after transfer of vitrified embryos. Thus, assuming that vitrification causes mitochondrial damages in the embryo, the resulting compromised functionality of these organelles could be inherited by later tissue cells [111-113]. Furthermore, proteomic results showed some downrepresented DEPs involved in the drug-metabolising mechanisms via cytochrome P450 in liver cells, reinforcing the compromised liver detoxification function insinuated by KEGG pathways of the transcriptomic comparison [114]. Finally, taking into account both omic studies in this work, it is interesting to highlight that 17 DETs and 32 DEPs are related to extracellular exosome GO term. Exosomes are membranous vesicles secreted by liver cells that contain proteins, lipids and nucleic acids coated with a lipid bilayer. Thus, the exosome load represents a snapshot of the parental cell metabolism at the time of release and has been proposed as a potential biomarker of liver disease [115]. So, this information reinforces the finding that the metabolism and physiology of hepatic tissue was modified after transfer of vitrified embryos. Proteomic analysis of these vesicles can offer interesting information to the field, increasing the studies that evaluate the ART effects through proteomic comparisons both in domestic animals [33-36], and humans [37, 116].

Hence, we noted that some global results seem to have a common denominator. Disturbed lipid metabolism and impaired mitochondrial function (energy metabolism) in the liver could be ascribed to early placental insufficiency that leads to foetal growth restriction [100, 117]. In this line, as we previously reported, evidence for placental abnormality in foetuses was observed after transfer of vitrified embryos [34, 35], probably due to preferential confinement of damaged cells to the trophectoderm [118]. In fact, it has been reported that gene expression related to lipid metabolism, steroidogenesis, cell differentiation and placentation changed in blastocyst embryos following cryopreservation [68]. In this sense, compromised placenta limits the availability of the critical substrates to the foetus and retards development of the embryo and/or its organs during gestation. This impaired organogenesis, particularly of the liver, could lead to permanent changes in glucose and lipid metabolism, accompanied by a disturbance in the oxidation of these substrates via *OXPHO*, which can affect the health of the offspring and could continue into later developmental stages until adulthood [18, 98-100, 117, 119]. Therefore, disruption in normal development may result in organogenic errors that could incur permanent changes observable in adult life. In concordance, liver but also heart weight was lower after embryo vitrification. Similar phenotype modifications were described in mice after *in vitro* culture without serum in the culture medium [7]. Accordingly, lower blood pressure was exhibited by IVF mice [13]. Notably, structural remodelling of the heart was exhibited by IVF children compared to spontaneously conceived offspring [120]. Transcriptomic analysis of the IVF progeny heart tissue revealed 1361 downregulated genes [38], suggesting that both heart structure and its physiology became modified after ART. Worryingly, these differences are generally only found after a careful post-mortem examination of apparently normal individuals, suggesting that a masked compromised welfare may be occurring [11].

## Conclusions

In conclusion, our experimental approach provides a broad overview that male progeny derived from vitrified embryo transfer manipulation technique have long-term consequences on growth rate and vital organ weights in adulthood, correlated with molecular signatures at transcriptomic and proteomic level. Today, a well-accepted hypothesis is that exposure of an organism to its environment at critical stages during development can trigger adaptive mechanisms, due to the active phenotypic plasticity of the embryo, resulting in a phenotypic variant of the individuals. This study should represent a significant step towards promoting a paradigm shift in characterisation of long-term consequences of ART in adulthood, and thus opens the way to elucidating the adaptive mechanisms of embryos from a systems biology perspective.

## Acknowledgements

English text version revised by N. Macowan English Language Service.

